# The Novel Monocomponent FAD-dependent Monooxygenase HpaM Catalyzes the 2-Decarboxylative Hydroxylation of 5-Hydroxypicolinic Acid in *Alcaligenes faecalis* JQ135

**DOI:** 10.1101/171595

**Authors:** Jiguo Qiu, Bin Liu, Lingling Zhao, Yanting Zhang, Dan Cheng, Xin Yan, Jiandong Jiang, Qing Hong, Jian He

## Abstract

5-hydroxypicolinic acid (5HPA) is a natural pyridine derivative that can be microbially degraded. However, the physiological, biochemical, and genetic foundation of the microbial catabolism of 5HPA remains unknown. In this study, a gene cluster *hpa* (which is involved in degradation of 5HPA in *Alcaligenes faecalis* JQ135) was cloned and HpaM was identified as a novel monocomponent FAD-dependent monooxygenase. HpaM shared a sequence only 31% similarity with the most related protein 6-hydroxynicotinate 3-monooxygenase (NicC) of *Pseudomonas putida* KT2440. *hpaM* was heterologously expressed in *E. coli* BL21(DE3), and the recombinant HpaM was purified via Ni-affinity chromatography. HpaM catalyzed the 2-decarboxylative hydroxylation of 5-HPA, thus generating 2,5-dihydroxypyridine (2,5-DPH). Monooxygenase activity was only detected in the presence of FAD and NADH, but not of FMN and NADPH. The apparent *K*_m_ values of HpaM toward 5HPA and NADH were 45.4 μ and 37.8 μ, respectively. Results of gene deletion and complementation showed that *hpaM* was essential for 5HPA degradation in *Alcaligenes faecalis* JQ135.

**Importance:** Pyridine derivatives are ubiquitous in nature and important chemical materials that are currently widely used in agriculture, pharmaceutical, and chemical industries. Thus, the microbial degradation and transformation mechanisms of pyridine derivatives received considerable attention. Decarboxylative hydroxylation was an important degradation process in pyridine derivatives, and previously reported decarboxylative hydroxylations happened in the C3 of the pyridine ring. In this study, we cloned the gene cluster *hpa*, which is responsible for 5HPA degradation in *Alcaligenes faecalis* JQ135, thus identifying a novel monocomponent FAD-dependent monooxygenase HpaM. Unlike 3-decarboxylative monooxygenases, HpaM catalyzed decarboxylative hydroxylation in the C2 of the pyridine ring in 5-hydroxypicolinic acid. These findings deepen our understanding of the molecular mechanism of microbial degradation of pyridine derivatives. Furthermore, HpaM offers potential for applications to transform useful pyridine derivatives.

## Introduction

Pyridine derivatives are common natural products, as well as important artificial compounds that are widely used in agriculture, pharmaceutical, and chemical industries as solvents, dyes, pharmaceuticals, herbicides, and pesticides (1-3). However, the increasing use of pyridine derivatives causes large amounts entering the environment, thus leading to severe environmental problems (4, 5). Therefore, the biodegradation or detoxication of pyridine derivatives, and their transformation to useful products are of significant interest.

Pyridine derivatives could either be degraded or transformed by a variety of bacteria, and the degradation processes are typically initialed via hydroxylation (6, 7). Nicotinic acid (NA, 3-pyridinecarboxylic acid) and nicotine often served as models for explore the catabolic mechanisms of pyridine derivatives (8-11). NA was initially hydroxylated at the C2 of the pyridine ring via NA monooxygenase (NicAB), thus producing 6-hydroxynicotinic acid (6HNA) (8). 6HNA was further decarboxylatively hydroxylated at the C3 of the pyridine ring via 6HNA monooxygenase (NicC) yielding 2,5-dihydroxypyridine (2,5-DHP), which was then subjected to ring-cleavage. Nicotine could be degraded via both the pyridine and pyrrolidine pathways, and both pathways include two hydroxylation steps. In the pyrrolidine pathway, the intermediate 3-succinoylpyridine (SP) was hydroxylated at the C2 of the pyridine ring via SP monoxygenase (Spm), thus generating 6-hydroxy-3-succinoylpyridine (HSP), which was further 3-decarboxylatively hydroxylated to 2,5-DHP via HSP monoxygenase (HspB). In the pyridine pathway, nicotine was hydroxylated at the C6 of the pyridine ring via nicotine hydroxylase (NDH) to 6-hydroxynicotine, while the downstream intermediate 2,6-dihydroxypyridine was 3-hydroxylated to 2,3,6-trihydroxypyridine via 2,6-dihydroxypyridine 3-monoxygenase (DHPH) (10, 12). NicAB and Spm are multicomponent molybdenum-containing monooxygenases, while the NicC and HspB are monocomponent flavin-dependent monooxygenases, catalyzing the 3-decarboxylatively hydroxylation. However, gene coding of monooxygenase catalyzing the 2-decarboxylatively hydroxylation of pyridine derivatives has not been reported to date.

5-Hydroxypicolinic acid (5HPA) is a isomer of 6HNA and a natural pyridine derivative produced by bacteria (such as *Nocardia* sp.) or plants (such as *Gynura divaricata*) (13, 14). The degradation of 5HPA has only been reported in *Pusillimonas* sp. 5HP (15). A 5-hydroxypicolinate 2-monooxygenase (catalyzing the 2-decarboxylative hydroxylation of 5-HPA to 2,5-DHP) was partially purified from strain 5HP. However, both the amino acid sequence of the 5-hydroxypicolinate 2-monooxygenase, and the genetic foundation of 5HPA degradation remain unknown.

In this study, the gene cluster *hpa* involved in 5HPA degradation was cloned, and a 5-<u>h</u>ydroxy<u>p</u>icolinic <u>a</u>cid 2-<u>m</u>onooxygenase HpaM was identified from *Alcaligenes faecalis* JQ135 (Fig. 1A, B). HpaM is FAD and NADH-dependent, and catalyzes the 2-decarboxylative hydroxylation of the pyridine-ring in 5HPA to produce 2,5-DHP.

**Fig. 1.**
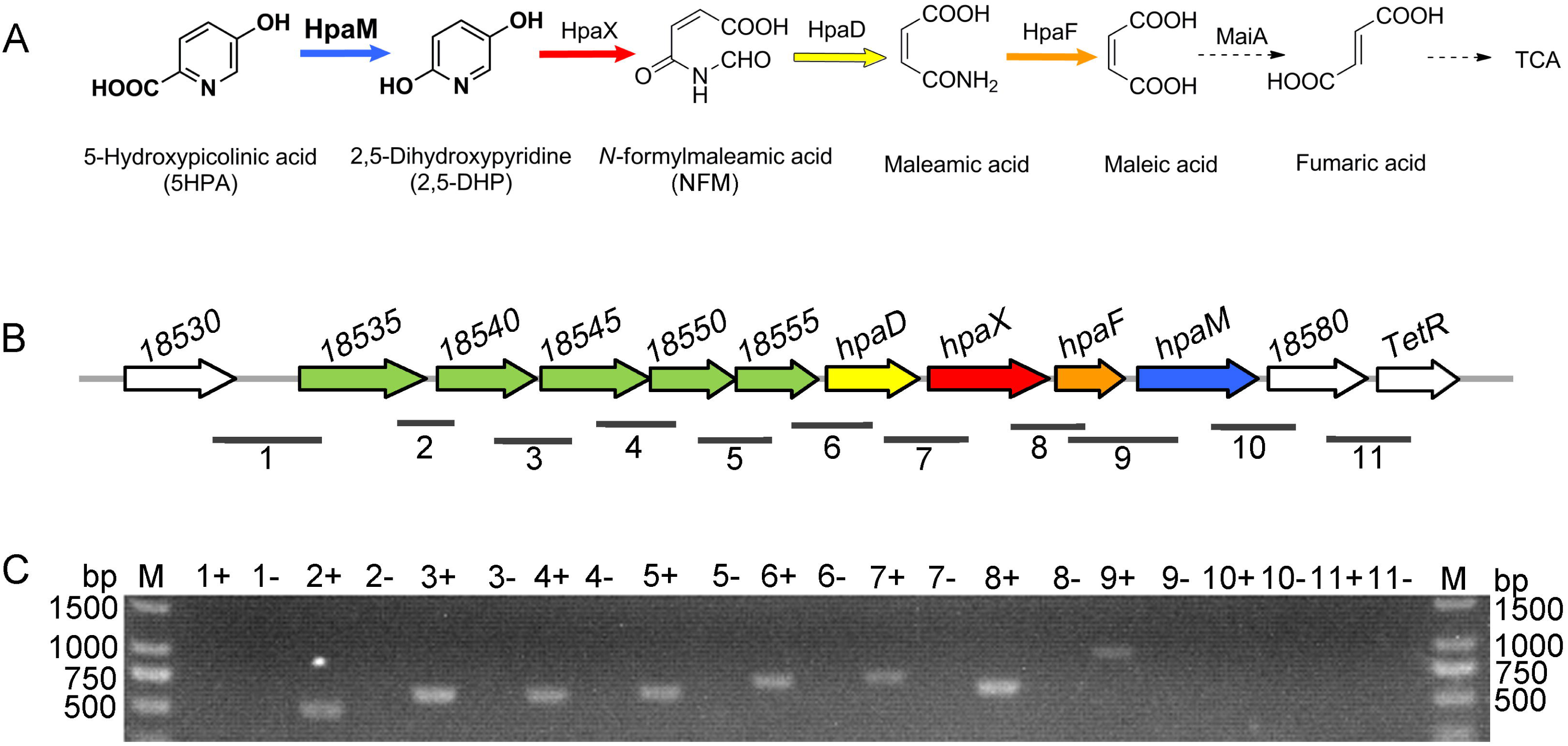
Catabolic mechanism of 5HPA in *A. faecalis* JQ135. (A) The proposed 5HPA degradation pathway in *A. faecalis* JQ135. HpaM, 5-hydroxypicolinic acid 2-monooxygenase; HpaX, 2,5-Dihydroxypyridine 5,6-dioxygenase; HpaD, *N*-formylmaleamic acid deformylase; HpaF, maleamic acid amidohydrolase; MaiA, maleic acid *cis-trans* isomerase; and TCA: tricarboxlic acid cycle. (B) Organization of the gene cluster *hpa*. *18530* indicates the locus tag of *AFA_18530*. Genes have been functionally annotated following the color code indicated in panel A. The lines below the gene cluster show the location and size of the PCR fragments in panel C. (C) Agarose gel electrophoresis of RT-PCR products. + and - indicate that the cells were cultured with 5HPA and glycerol, respectively.

## Results

### Degradation of 5HPA by strain *A. faecalis* JQ135

*A. faecalis* JQ135 was formerly identified as a picolinic acid (PA)-degrading bacterium (16). 5HPA is a 5-hydroxylated derivate of PA that was tested for degradation and utilization by the strain in a carbon and nitrogen-absent MSM. The results showed that the strain *A. faecalis* JQ135 could completely degrade 1 mM 5HPA within 36 h, and correspondingly, the OD600 of the culture increased from 0.2 to 0.5 ± 0.1 (Fig. 2). These results indicated that strain *A. faecalis* JQ135 could degrade and utilize 5HPA as sole source of carbon and nitrogen for growth. In addition, attempts to detect metabolic intermediates of 5HPA within the culture failed. This might because little or no intermediates were excreted from strain JQ135 cells during degradation of 5HPA.

**Fig. 2.**
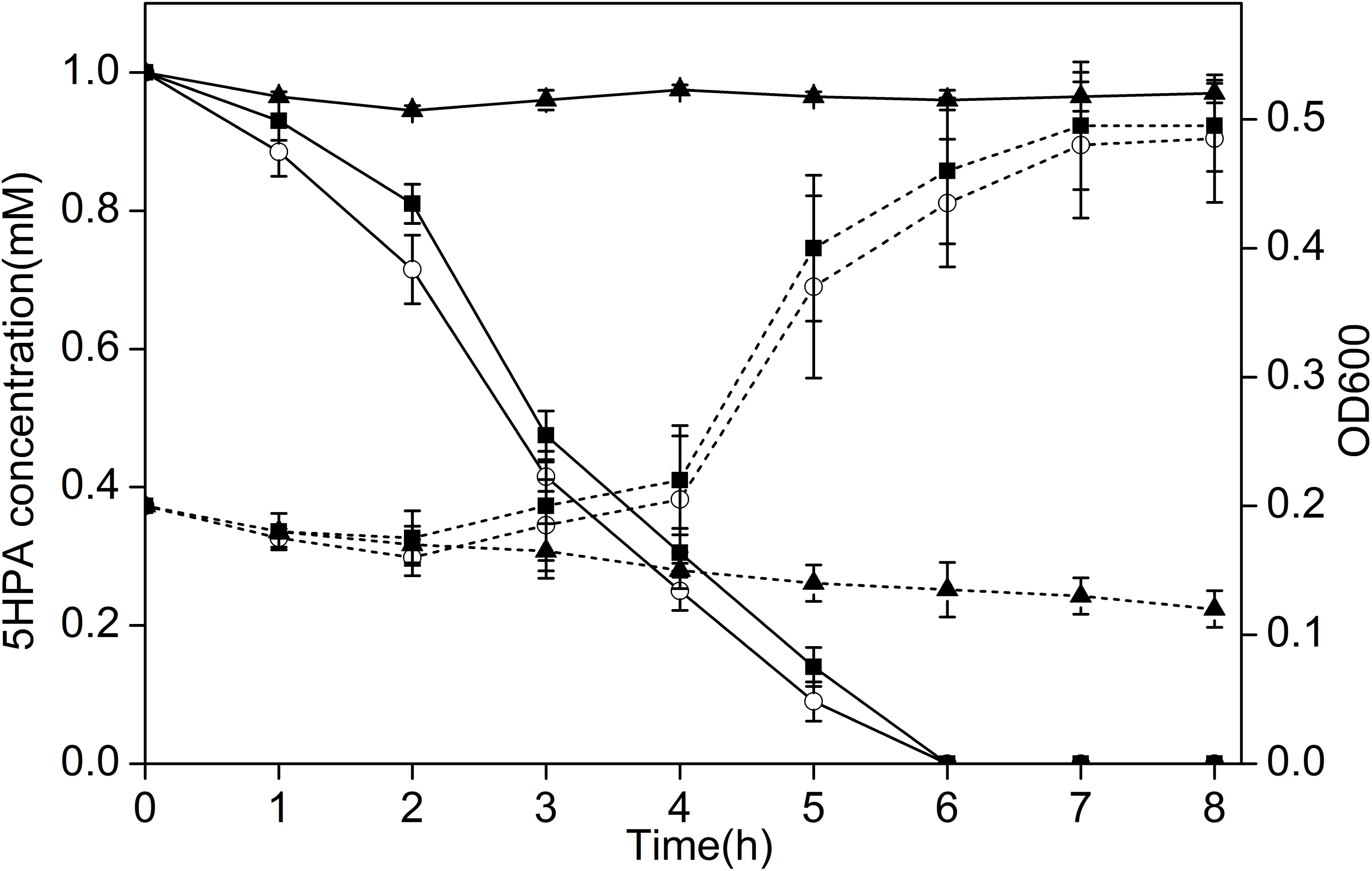
Degradation of 5HPA (solid lines) and cell growth (dotted lines) of the wild-type strain JQ135 (▪), the mutant Z10 (▲), and the complemented strain Z10-pBBR-*hpaM* (○).

### Transposon mutagenesis and cloning of a gene cluster involved in 5HPA degradation

To clone the genes involved in 5HPA degradation, a transposon mutagenesis library of *A. faecalis* JQ135 was constructed. One mutant (Z10) could not grow in MSM agar containing 1 mM 5HPA and was screened from approximately 5000 mutants. When inoculated into liquid MSM, containing 1 mM 5HPA, mutant Z10 could not degrade 5HPA (Fig. 2). Furthermore, the genome of *A. faecalis* JQ135 was determined by the PacBio system. The complete genome of the strain contained one circular chromosome (4,078,346 bp) and no plasmid could be found. A total of 3,723 ORFs were predicted. The insertion position of the transposon, determined via the DNA walking method (17), was located in gene *AFA_18575* (genome position 4,070,825). *AFA_18575* is 1,218 bp in length with a G+C content of 56.08%. The deduced protein was searched against the NCBI database (http://blast.ncbi.nlm.nih.gov/), using the BLASTP program (Table 1). The results showed that the proteins that were related the most were flavin-containing monooxygenases, such as 6-hydroxynicotinate 3-monooxygenase (NicC, sequence ID: Q88FY2) from *Pseudomonas putida* KT2440 (identity of 31%) (8), and salicylate 1-monooxygenase (SalM or NahG, sequence ID: P23262) from *P. putida* KF715 (identity of 28%) (18) (Fig. 3, S1). NicC and SalM catalyzed the decarboxylatively hydroxylation of 6HNA and salicylate, respectively. Downstream of *AFA_18575*, a gene (*AFA_18580*) with unknown function in the DUF2236 family and a TetR-type regulator gene (*AFA_18585*) followed (Fig. 1B; Table 1). Upstream of *AFA_18575*, eight genes were found (*AFA_18535* to *AFA_18570*). AFA_18535, AFA_18540, and AFA_18545 were predicted to be branched-chain amino acid ABC transporter substrate-binding proteins, while AFA_18550 and AFA_18555 were branched-chain amino acid ABC transporter ATP-binding proteins, indicating that these five genes encoded an ABC-type transporter. The next three genes *AFA_18560*, *AFA_18565*, and *AFA_18570*, showed 57%, 53%, and 41% identities, respectively, with NicD (*N*-formylmaleamaic acid deformylase), NicX (2,5-DHP dioxygenase), and NicF (maleamate amodohydrolase) from *P. putida* KT2440 at amino acid levels (Fig. 3). Interestingly, NicD, NicX, and NicF, in combination with NicC, formed a complete metabolic pathway for the transformation of 6HNA into maleic acid (8). Considering the structural similarity between 5HPA and 6HNA, we presumed that these contiguous genes were organized in a single cluster as well as involved in the degradation of 5HPA in *A. faecalis* JQ135. For this study, this gene cluster was designated as *hpa* (<u>h</u>ydroxy<u>p</u>icolinic <u>a</u>cid).

**Table 1.**
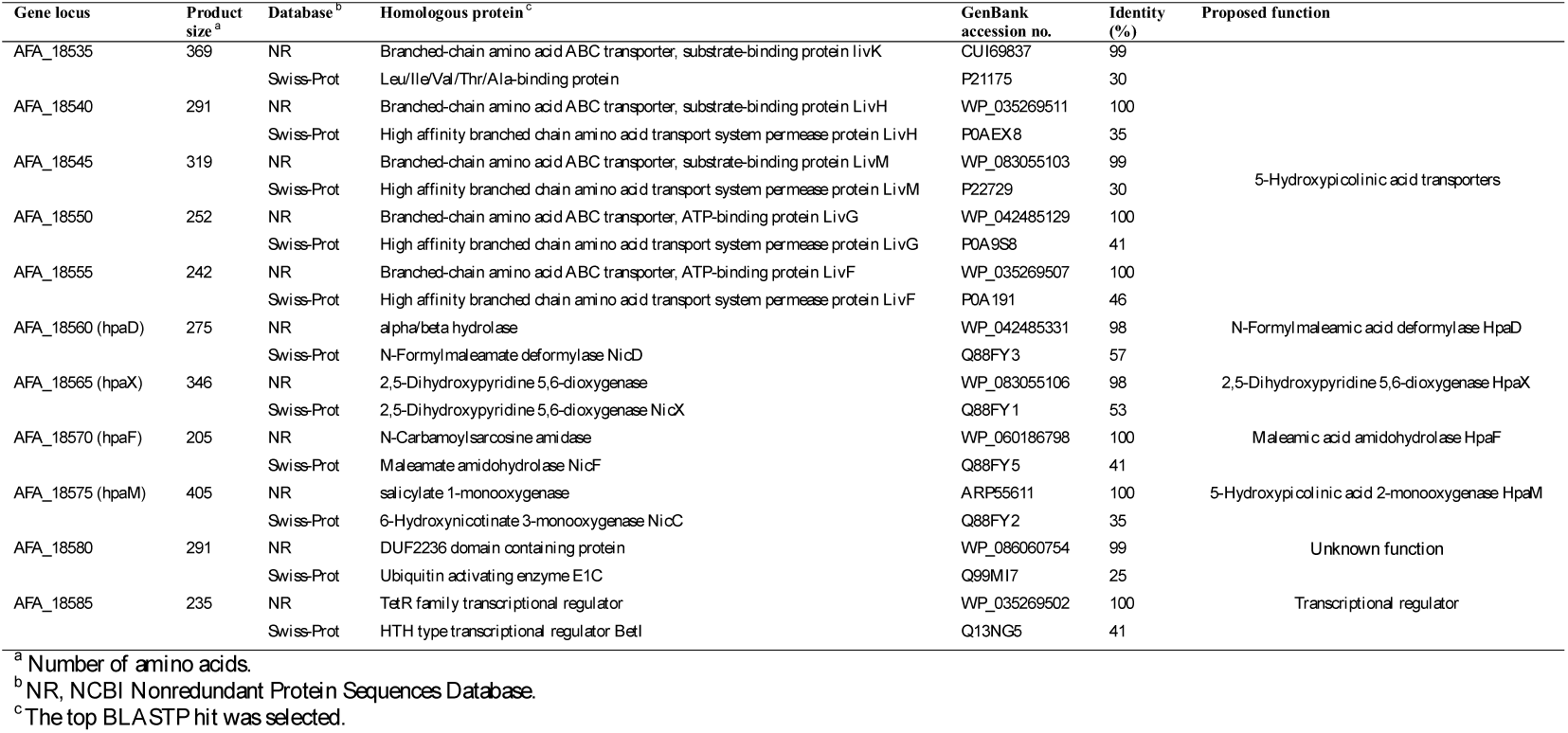
Sequence comparisons of *hpa* gene cluster of *A. faecalis* JQ135 with database entries.

**Fig. 3.**
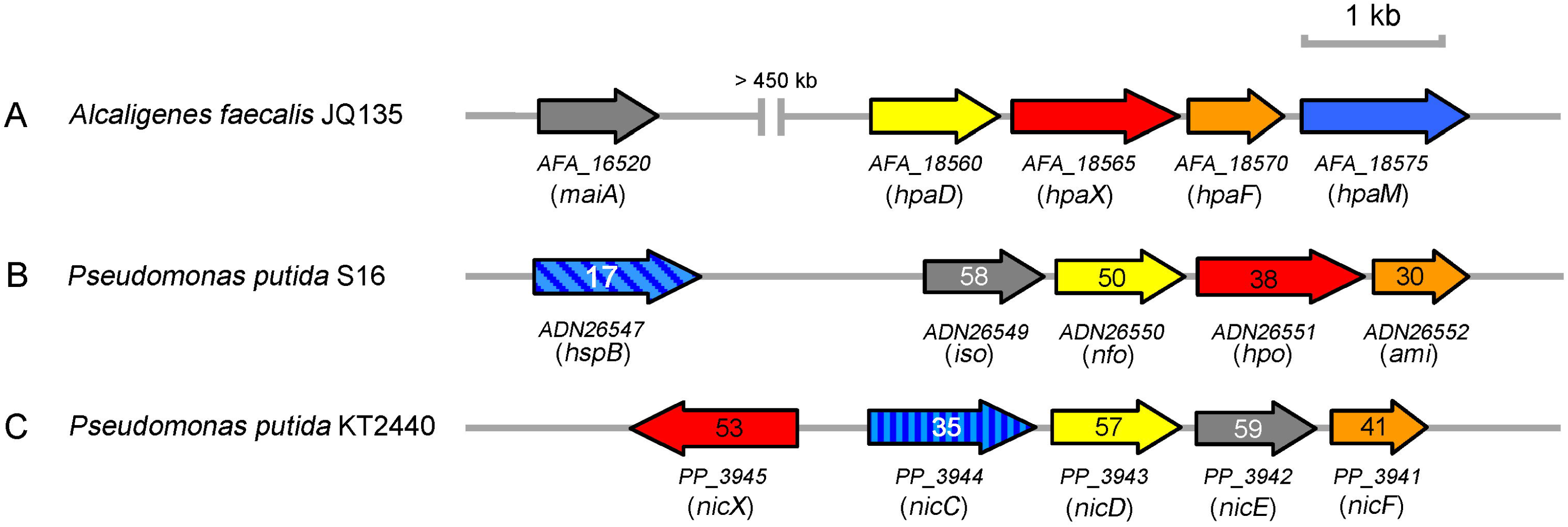
Organization of the *hpa* cluster for 5HPA degradation in *A. faecalis* JQ135 (A), the *nic2* cluster for nicotine degradation in *P. putida* S16 (B), and the *nic* cluster for nicotinic acid degradation in *P. putida* KT2440 (C) (8). *AFA_16520* (*maiA*): locus tag (gene name). Numbers within arrows indicate the protein sequence identity with the orthologous from strain JQ135. *hpaM*: 5-hydroxypicolinic acid 2-monooxygenase gene; *hspB*: 6-hydroxy-3-succinoylpyridine 3-monooxygenase gene; *nicC*: 6-hydroxynicotinate 3-monooxygenase gene; *hpaX*, *hpo*, and *nicX*: 2,5-dihydroxypyridine 5,6-dioxygenase gene; *hpaD*, *nfo*, and *nicD*: *N*-formylmaleamate deformylase gene. *hpaF*, *ami*, and *nicF*: maleamate amidohydrolase; *maiA*, *iso*, and *nicE*: maleic acid *cis-trans* isomerase gene.

To confirm the function of *hpaM* (*AFA_18575*), a fragment containing *AFA_18575* was amplified from the genome of *A. faecalis* JQ135 and ligated into the broad-host-range vector pBBR1MCS-5, thus generating pBBR-*hpaM.* The mutant Z10, which was introduced with pBBR-*hpaM* (Z10-pBBR-*hpaM*) regained the ability to degrade 5HPA (Fig. 2), suggesting that HpaM enabled the conversion of 5HPA.

### Gene knockout of *hpaM* and the phenotype analysis

To further determine the physiological function of *hpaM*, both the in-frame deletion mutant JQ135Δ*hpaM* and complemented strain JQ135Δ*hpaM*-pBBR-*hpaM* were constructed. The strain JQ135Δ*hpaM* had lost the ability to degrade and utilize 5HPA, while the complemented strain JQ135Δ*hpaM*-pBBR-*hpaM* recovered the ability to degrade and grow on 5HPA. These results clearly demonstrate the gene *hpaM* to be essential for the catabolism of 5HPA in *A. faecalis* JQ135.

### Transcription levels of genes in cluster *hpa* in response to 5HPA

Real-time PCR (RT-PCR) was conducted to determine the transcription of the *hpa* cluster and to investigate whether these genes could be induced by 5HPA. The results showed that the five putative transporter-coding genes (*AFA_18535* to *AFA_18555*), *hpaD*, *hpaX*, *hpaF*, and *hpaM* comprised a transcriptional operon, and were transcribed in cells grown on 5HPA containing medium, but not on glycerol (Fig. 1C), indicating that the cluster *hpa* was induced by 5HPA. These results further indicate that cluster *hpa* was involved in the degradation of 5HPA in *A. faecalis* JQ135.

### Heterogenous expression and purification of HpaM and function determination

*hpaM* was cloned into pET29a(+) and expressed in *E. coli* BL21(DE3). The N-terminal 6×His-tag HpaM was purified via nickel affinity chromatography. The purified enzyme migrated as a single band with the size of about 45 kDa (obtained via SDS-PAGE analysis), which was in agreement with its theoretical value (Fig. S2).

HpaM was predicted to contain FAD binding domains (such as GXGXXG) (Fig. S1). However, the purified HpaM remained colorless, suggesting that no flavin was associated after purification. The purified HpaM showed enzymatic activity only after addition of external FAD (but not FMN). NADH (but not NADPH) could be used as electron donor. These results indicated that HpaM was FAD- and NADH-dependent. The rapid degradation of 5HPA was monitored via spectrophotometric changes (250 nm to 400 nm). First, the absorption spectra of authentic 5HPA (maximum UV absorption (λ_max_) at 278 and 320 nm), 2,5-DHP (λ_max_ at 322 nm), and NADH (λ_max_ at 272 and 340 nm) in 50 mM PBS (pH7.0) were determined (Fig. 4B). Then, the reaction was started with addition of 5HPA, and the consumptions of 5HPA (at 278 nm) and NADH (between 340 to 400 nm) were observed (Fig. 4C). HPLC results showed that 5HPA (with a retention time of 5.52 min) decreased and a product (with a retention time of 5.22 min) accumulated (Fig. S3). The retention time of the compound was equal to that of the authentic 2,5-DHP standard, and the LC/MS analysis indicated that the product showed a molecular ion at *m/z* 112.1 [M+H]^+^, which was identical to that of 2,5-DHP in previous reports (12, 19). Thus, based on the above analysis, the product could be identified as 2,5-DHP.

**Fig. 4.**
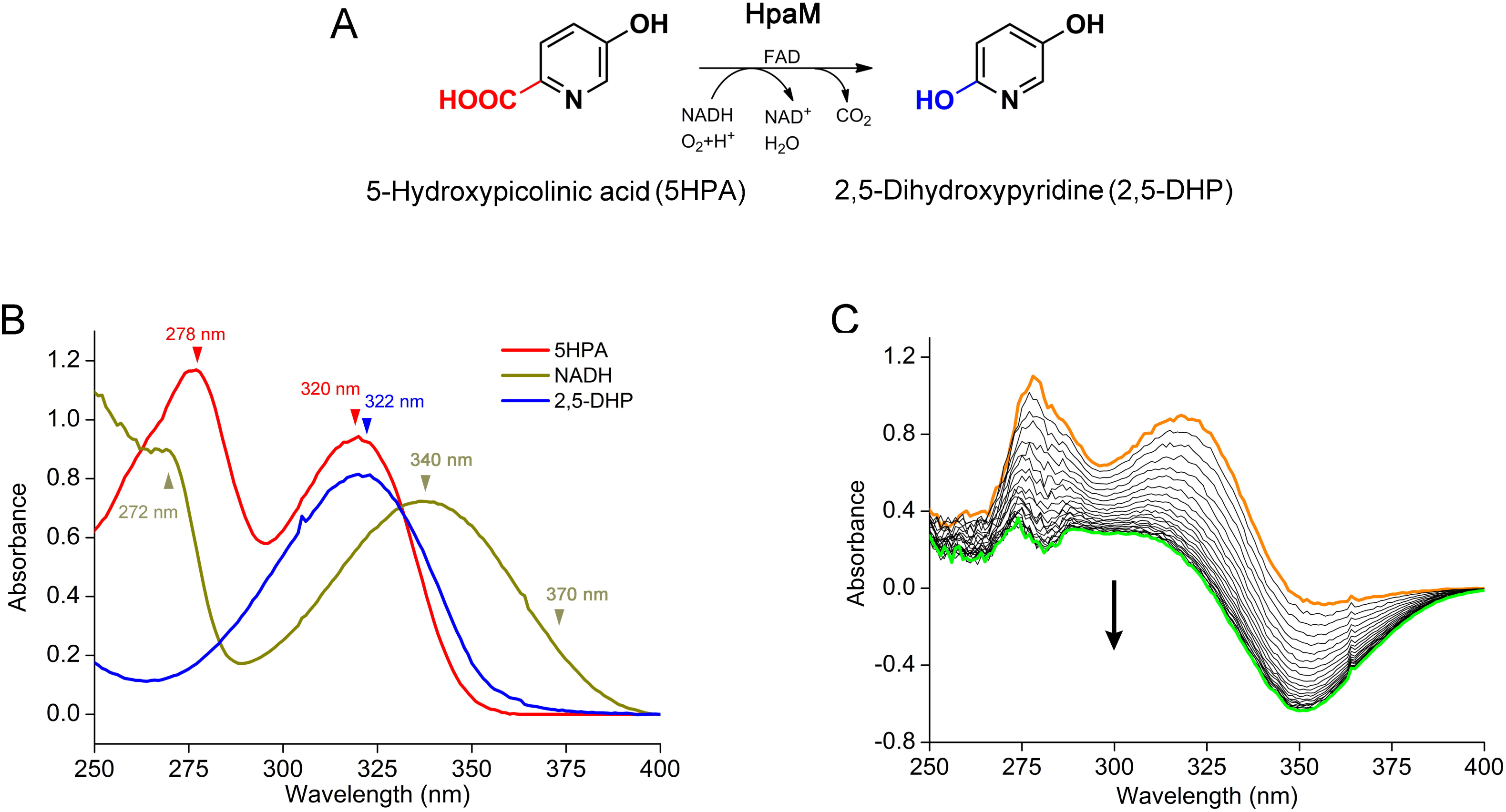
Rapid determination of the HpaM activity. (A) Proposed conversion of 5HPA to 2,5-DHP catalyzed via HpaM. (B) UV absorption spectrum of authentic 5HPA, 2,5-DHP, and NADH in 50 mM PBS (at pH 7.0). (C) Spectrophotometric changes during the transformation of 5HPA via HpaM. The reference cuvette contained HpaM, FAD, and NADH. The reaction was initiated via addition of 5HPA. The spectra were recorded every 90 s. The arrow indicates the direction of spectral changes.

### Kinetics analysis constants and biochemical properties

Kinetic analysis revealed *K*_m_ and *k*_cat_ values of HpaM for 5HPA of 45.4 ± 4.2 μ and 10.2 ± 0.3 s^-1^, respectively (Fig. S4). The apparent catalytic efficiency (*k*_cat_*/K*_m_) was 225.5 s^-1^ mM^-1^. Additionally, the apparent *K*_m_ value of HpaM for NADH was 37.8 ± 3.4 μM (Fig. S4). HpaM showed activities within a narrow pH range (pH 6.0 to 8.0) (Fig. S5). The optimal pH was found to be 7.0 in 50 mM PBS. Most of the activity was lost at pH 3.0 and 9.0. The optimal temperature was 25°C. It retained approximately 60% of its activity at 20°C and 35°C. HpaM remained stable for seven days at 4°C in PBS (50 mM, pH 7.0). As the temperature increased, the enzyme was unstable. It lost 60% of the activities after incubation in 40°C for 1 h. When 1 mM of various metal ions were added into the reaction mixture, the Ag^+^, Cu2^+^, Hg^2+^, Ni^2+^, or Zn^2+^ completely inhibited the activities, while only a slight inhibition was found for Ca^2+^, Co^2+^, Fe^3+^, Li^+^, or Mg^2+^, and slight activation was found for Mn^2+^.

The following structural analogs of 5HPA were tested as substrates in a standard activity test: PA, NA, 5-aminopicolinic acid, 5-methylpicolinic acid, 5-chloropicolinic acid, 5-bromoopicolinic acid, 3-hydroxypicolinic acid, 6-hydroxypicolinic acid, 2,3-pyridinedicarboxylic acid, 2,5-pyridinedicarboxylic acid, 2,6-pyridinedicarboxylic acid, 2-hydroxynicotinic acid, 4-hydroxynicotinic acid, 5-hydroxynicotinic acid, 6-hydroxynicotinic acid, and 3-hydroxyisonicotinic. HpaM showed no activities toward any of these compounds, indicating that the hydroxylase activity was 5HPA specific.

## Discussion

In this study, the 5HPA-degradation-deficient mutant JQ135m was screened from the Tn5-transposon mutant library of *A. faecalis* JQ135. DNA walking results and bioinformatics analysis revealed that the transposon was inserted into the monoxygenase gene *hpaM*. HpaM was shown to be responsible for the initial 2-decarboxylative hydroxylation of 5HPA based on the following indications, 1) deletion of *hpaM* resulted in a deficient 5HPA degradation in *A. faecalis* JQ135Δ*hpaM*, while introduction of *hpaM* into JQ135Δ*hpaM* regained the ability to degrade 5HPA; 2) the transcription of the *hpaM* was evidently induced by 5HPA; and 3) purified HpaM could transform 5HPA to 2,5-DPH *in vitro*.

Hydroxylation increased the hydrophilicity and polarity of substrates, thus this is typically the initial and key degradation step of aromatic compounds. So far, a number of flavin-dependent monooxygenases have been described that are responsible for the hydroxylation of aromatic compounds. Most of them catalyzed the direct hydroxylation of unsubstituted carbon atoms within the aromatic rings, and there were also a few flavin-dependent monooxygenases that catalyzed the decarboxylative hydroxylation of carboxyl-substituted carbon atoms in the aromatic ring (Fig. 5C). So far, only two of them (NicC and HspB) catalyzed the 3-decarboxylative hydroxylation of the pyridine-ring (Fig. 5B) (8, 20, 21). To the best of our knowledge, HpaM was the first identified flavin-dependent monooxygenase that catalyzed the 2-decarboxylative hydroxylation of pyridine or pyridine derivatives. Our report provides a novel insight into the decarboxylative 2-hydroxylation of pyridine derivatives.

**Fig. 5.**
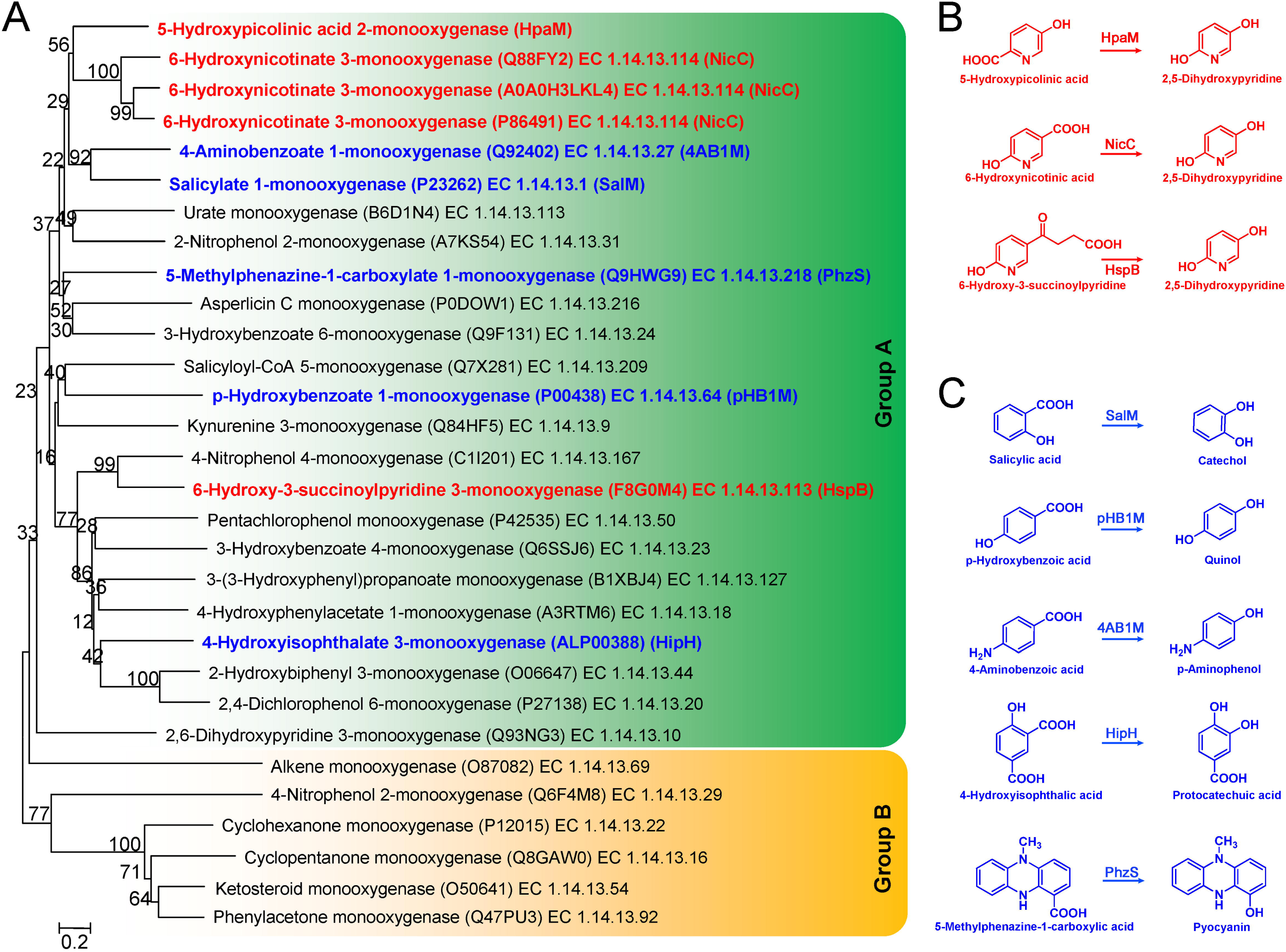
Classification analysis of HpaM and several related flavin-dependent monooxygenases. (A) Phylogenetic tree construction based on the alignment of HpaM with related monocomponent flavin-dependent monooxygenases in Group A (which contained one Rossmann fold) and Group B (which contained two Rossmann folds) (22, 23). Multiple-alignment analysis was performed via ClustalX v2.0. The phylogenetic tree was constructed via the neighbor-joining algorithm using MEGA 6.0 and bootstrap values (based on 1,000 replications) have been indicated at branch nodes. Bar, 0.20 substitutions per nucleotide position. Each item was arranged in the following order: protein name, UniProtKB/SwissProt accession number, and EC number. (B) Reactions of HpaM and two monooxygenases, catalyzing the 3-decarboxylative hydroxylation of pyridine derivatives. NicC, 6-hydroxynicotinate 3-monooxygenase (EC 1.14.13.114); HspB, 6-hydroxy-3-succinoylpyridine 3-monooxygenase (EC 1.14.13.163). (C) Five decarboxylative hydroxylation reactions of the benzene ring. SalM, salicylate 1-monooxygenase (EC 1.14.13.1); pHB1M, *p*-hydroxybenzoate 1-monooxygenase (EC 1.14.13.64); 4AB1M, 4-aminobenzoate 1-monooxygenase (EC 1.14.13.27); HipH, 4-Hydroxyisophthalic acid 3-monooxygenase; PhzS, 5-methylphenazine-1-carboxylate 1-monooxygenase (EC 1.14.13.218). The proteins in panels B and C of the phylogenetic tree have been indicated in red and blue, respectively.

HpaM showed low similarities (identities of only 31-28%) to several flavin-dependent monooxygenases. Flavin-dependent monooxygenase, which contains the vitamin B2 derivatives FAD and FMN as redox-active prosthetic group, is the largest family of flavoenzymes. To date, more than 230 flavin-dependent monooxygenases have been registered in class 1.14.13.- (http://enzyme.expasy.org/EC/1.14.13.-). These catalyze a variety of reactions such as, hydroxylation, Baeyer–Villiger oxidation, sulfoxidation, epoxidation, and halogenation, while playing an important role in the catabolism of natural and xenobiotic compounds (22). They are divided into six groups (A to F), and groups A and B are monocomponent systems, while groups C–F are two-component systems. Group A flavin monooxygenases rely on NAD(P)H as external electron donor and contain a glutathione reductase (GR-2) type Rossmann fold (GXGXXG) for FAD binding (22, 23). Group B flavin monooxygenases are similar to group A except that they contain two Rossmann folds. Sequences alignment showed that HpaM contained several conserved motifs, such as GXGXXG, DGX_5_R, and GDAX_10_GX_6_DX_3_L (Fig. S1) (23, 24). On the phylogenetic tree, which was constructed based on related monocomponent flavin-dependent monooxygenases, HpaM clearly located within Group A, forming a subclade with 6-hydroxynicotinate 3-monooxygenase NicC (Fig. 5A). The evidence indicates that HpaM is a novel group of A flavin monooxygenases.

Bioinformatic analysis and RT-PCR results indicated that *hpaM* and its upstream eight genes (*AFA_18535* to *AFA_18570*) constituted a 5HPA-inducible transcriptional operon. *AFA_18535* to *AFA_18555* were predicted to encode the five subunits of an ABC-type transporter, which may be responsible for the transmembrane transport of 5HPA. AFA_18560 (*hpaD*), AFA_18565 (*hpaX*), and AFA_18570 (*hpaF*) showed the highest similarities with 2,5-DHP catabolic enzymes NicD, NicX, and NicF from *P. putida* KT2440, respectively (Fig. 3 and Table 1). This suggests that the genes in cluster *hpa* constituted a catabolic pathway of 5HPA. However, it is interesting that the cluster *hpa* lacked a maleate isomerase gene, which is essential for the transformation of maleic acid to fumaric acid, while this maleate isomerase gene (*nicE* or *iso*) is present in the 2,5-DHP metabolic gene cluster in *P. putida* strains KT2440 and S16 (8, 25). However, unpublished data showed that strain JQ135 could utilize maleic acid as the sole carbon source for growth. Bioinformatics analysis showed that a gene (*AFA_16520*), which was physically separated from the *hpa* cluster, showed 99% identity with a previously reported maleic acid *cis-trans* isomerase (MaiA) from *A. faecalis* IFO13111 and 59% identity to NicE from *P. putida* KT2440 at the amino acid level (26). MaiA catalyzed the reversible conversion of maleate to fumarate. Thus *AFA_16520* was predicted to convert maleate to fumarate in *A. faecalis* JQ135. Based on the above analysis, we proposed a 5HPA degradation pathway in *A. faecalis* JQ135: (a) 5HPA is 2-decarboxylative hydroxylated by HpaM, generating 2,5-DHP; (b) 2,5-DHP is further ring-cleaved into *N*-formylmaleamic acid (NFM); (c) NFM is subsequently transformed to fumaric acid (a metabolite of citric acid cycle) via maleamic acid and maleic acid as intermediates (Fig. 1). Besides 5HPA, *A. faecalis* JQ135 could also utilize PA, NA, and 6-hydroxypicolinic acid. Maleic acid was the common intermediate metabolite of all above compounds, thus *AFA_16520* might be involved in the mineralization of all these compounds. The separation of *AFA_16520* from the *hpa* cluster might facilitate the flexible regulation of *AFA_16520* for the degradation of these substrates in strain JQ135. Moreover, the homologous genes of *AFA_16520* and *hpa* cluster were highly conserved (> 95% identities at the level of amino acid sequences) in most (13 out of 16) *A. faecalis* strains with available genome in GenBank (https://www.ncbi.nlm.nih.gov/genome/genomes/13038?; Table S1), suggesting that the degradation of 5HPA was a common feature of *A. faecalis*.

Additionally, 2,5-DHP (the hydroxylated product of 5HPA) could easily be chemically transformed to 5-aminolevulinic acid, which is an important chemical intermediate used for synthesis of plant hormones and drugs in cancer diagnosis and therapy. Thus, our present study might provide a new biological approach to produce 2,5-DHP.

## Materials and Methods

### Chemicals and media

5HPA (99%) and its structural analogs were purchased from J&K Scientific Ltd. (Shanghai, China). 2,5-DHP (98%) was purchased from SynChem OHG (Altenburg, Germany). All other chemicals and solvents used in this experiment were commercially available. Enzymes used in this study were purchased from Vazyme Biotech Co., Ltd (Nanjing, China). Mineral salts medium (MSM) and Luria-Bertani medium (LB) have been described in previous report (16), and to prepare nitrogen-absent MSM, NH_4_NO_3_ was replaced.

### Bacterial strains, vectors, and growth conditions

Bacterial strains and vectors used in this study are listed in Table 2. *A. faecalis* JQ135 has previously been identified as a picolinic acid-degrading bacterium (CCTCC M 2015812) (16). *E. coli* strains were grown at 37°C, while other strains were grown at 30°C. The media were supplemented with chloramphenicol (Cm, 34 μg/mL), kanamycin (Km, 50 μg/mL), gentamicin (Gm, 50 μg/mL), or streptomycin (Str, 50 μg/mL) as required.

**Table 2.**
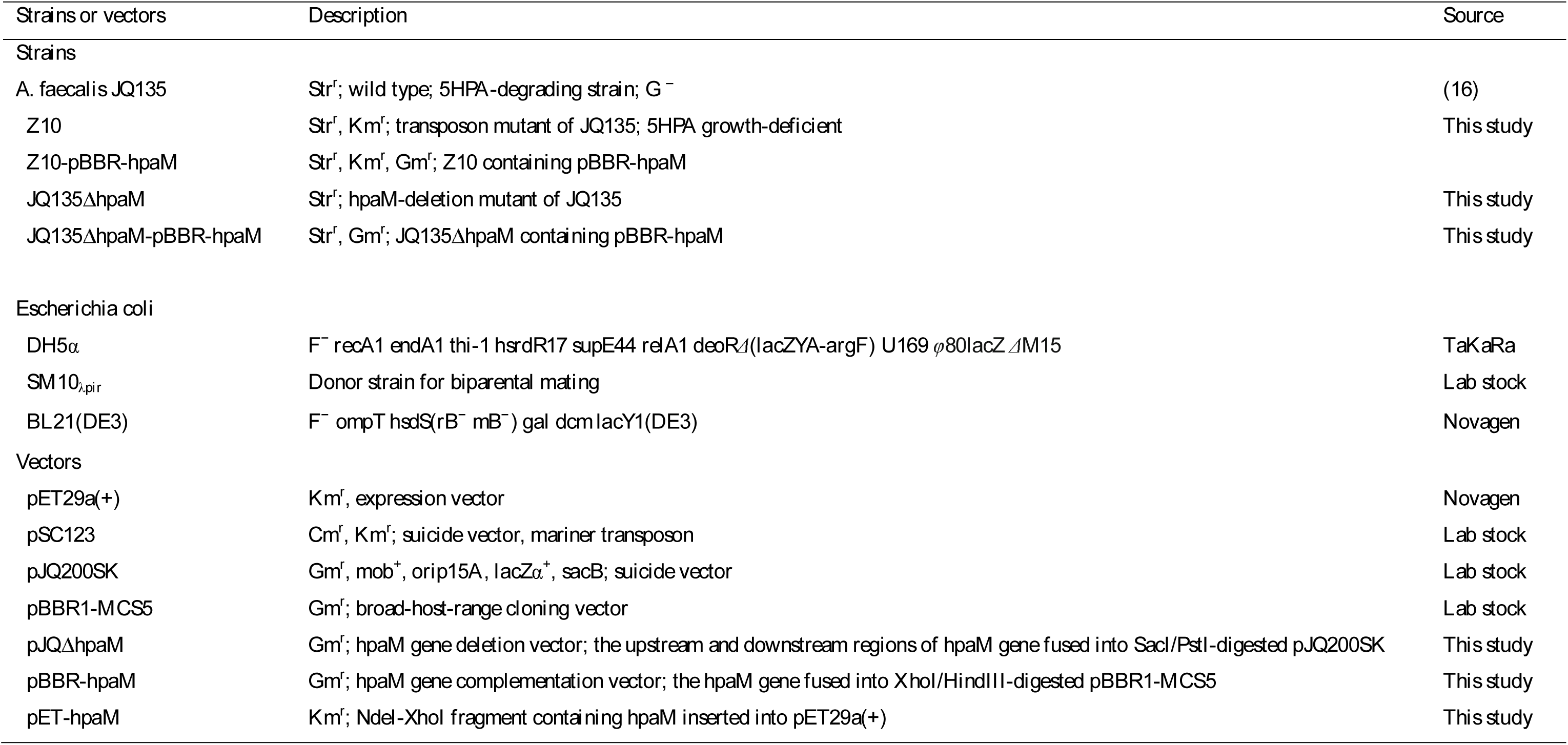
Strains and vectors used in this study.

### Isolation of 5HPA growth-deficient mutant and determination of the transposon insertion site

Both generation and selection of 5HPA growth-deficient mutants of *A. faecalis* JQ135 were performed according to the selection method via picolinic acid (PA) growth-deficient mutants as previously described (16). PA was replaced by 5HPA. The insertion site of the transposon was determined via SEFA-PCR as previously described (16, 17).

### Deletion of *hpaM* and complementation

Standard DNA manipulation was performed as previously described (27). All primers used in this study are listed in Table S2. The deletion of the *hpaM* gene in *A. faecalis* JQ135 was generated via a two-step homogenetic recombination method using the suicide vector pJQ200SK. The two primer pairs, koUF/koUR and koDF/koDR, were used to amplify the homologous recombination-directing sequences. Then, both PCR fragments were cloned into SacI/PstI-digested pJQ200SK using the ClonExpress MultiS One Step Cloning Kit (Vazyme Biotech Co., Ltd, Nanjing, China). The resulting vector pJQ-Δ*hpaM* was then introduced into *A. faecalis* JQ135 cells. The single-crossover mutant was screened on LB plates containing Str and Gm. The double-crossover mutant (JQ135Δ*hpaM*) was selected on LB plates mixed with Str and 10% (wt/vol) sucrose. The vector pBBR-*hpaM* was constructed for gene complementation. The *hpaM* gene was amplified via primers *hpaM*-F and *hpaM*-R, and then fused with the XhoI/HindIII-digested pBBR1-MCS5, thus generating pBBR-*hpaM*. The pBBR-*hpaM* vector was transferred into the mutant JQ135Δ*hpaM* via triparental mating to generate the complemented strain JQ135Δ*hpaM*-pBBR-*hpaM*.

### RNA extraction and RT-PCR

JQ135 Cells were cultured in glycerol and harvested at the mid-exponential phase, washed twice with MSM, and resuspended in MSM. The cell suspension (with an OD600 of 0.6) was transferred into a 50 mL flask, containing 20 mL MSM supplemented with 1 mM of either glycerol or 5HPA. After incubation at 30°C and 180 rpm for 6 h, cells were harvested. Total RNA was isolated using an RNA isolation kit (TaKaRa). Reverse transcription (RT)-PCR was conducted with a PrimeScript RT reagent kit (TaKaRa). All RT-PCR primers are listed in Table S2. All samples were run in triplicate.

### Cloning, overexpression, and purification of HpaM

The *hpaM* gene (ignoring the stop codon) was PCR amplified with primers exp*hpaM*F and exp*hpaM*R and fused into the *Nde*I/*Xho*I digested pET29a(+), thus producing pET-*hpaM*. The C-terminal 6×His-tagged HpaM was overexpressed in *E. coli* BL21(DE3) that carried pET-*hpaM*. The cells were grown in LB at 37°C to an OD600 of 0.5 and then induced for 16 h via addition of 0.1 mM isopropyl-β-D-thiogalactopyranoside (IPTG) at 16°C. HpaM was purified via Ni^2+^-nitrilotriacetic acid agarose chromatography (Novagen) and eluted at 100 mM imidazole. HpaM was dialyzed against PBS (50 mM, pH 7.0) at 4°C for 24 h and analyzed via 12.5% SDS-PAGE.

### Enzyme assays

The standard reaction mixture contained 50 mM PBS (pH 7.0), 0.2 mM FAD, 0.5 mM NADH, 0.2 mM 5HPA, and 1 μg purified HpaM. The reference cuvette contained all of these compounds except for 5HPA. The assay was initiated via addition of substrate 5HPA. For the activity assay, HpaM activity was spectrophotometrically analyzed by measuring NADH oxidation at 370 nm (ε = 2,470 M^-1^ cm^-1^) (28) instead of at its λ_max_ of 340 nm to avoid interference of the absorptions of 5HPA and 2,5-DHP (Fig. 4B). One unit of HpaM activity was defined as the amount of enzyme required for the oxidation of 1 μmol of NADH per min at 25°C. Rapid observation of HpaM activities was monitored via spectrophotometric changes from 250 nm to 450 nm at room temperature, using a UV2450 spectrophotometer (Shimazu) in 1 cm path length quartz cuvettes. For determination of kinetics constants, substrates were appropriately diluted into seven concentrations around the *K_m_* (3.6, 7.2, 18, 36, 72, 108, 144, and 360 μM for 5HPA; 5, 10, 25, 50, 75, 100, and 500 μM for NADH). The kinetics values were calculated via nonlinear regression fitting to the Michaelis-Menten equation. The effects of different pH (3.0 to 10.0), temperature (10 to 60°C), and metal ions on HpaM activities were determined as previously reported (29).

### Analytical methods

Determinations of 5HPA and 2,5-DHP were performed via HPLC analysis on an Shimadzu AD20 system equipped with Phecda C18 reversed phase column (250 mm × 4.60 mm, 5 μm) with array detection at both 280 nm and 310 nm. The mobile phase consisted of methanol : water : formic acid (12.5:87.5:0.2, v/v/v) at a flow rate of 0.6 mL/min, 30°C. All assays in this study were independently performed three times, and the means and standard errors of measurements were calculated. LC/MS analysis was performed in a Thermo (America) DECA-60000 XLCQ Deca XP Plus instrument as previously described (19).

### Nucleotide sequence accession numbers

The *hpa* cluster sequence and the complete genome sequence of *A. faecalis* JQ135 have been deposited in the GenBank database under accession numbers KY230187 and CP021641, respectively.

## Acknowledgments

This work was supported by the National Natural Science Foundation of China (No. 31500082), China Postdoctoral Science Foundation (No. 2016M601826), the Postdoctoral Foundation of Jiangsu Province (No. 1601035A), and Natural Science Foundation of Jiangsu Province (No. BK20141366).

